# ntJoin: Fast and lightweight assembly-guided scaffolding using minimizer graphs

**DOI:** 10.1101/2020.01.13.905240

**Authors:** Lauren Coombe, Vladimir Nikolić, Justin Chu, Inanc Birol, René L. Warren

## Abstract

**Summary:** The ability to generate high-quality genome sequences is cornerstone to modern biological research. Even with recent advancements in sequencing technologies, many genome assemblies are still not achieving reference-grade. Here, we introduce ntJoin, a tool that leverages structural synteny between a draft assembly and reference sequence(s) to contiguate and correct the former with respect to the latter. Instead of alignments, ntJoin uses a lightweight mapping approach based on a graph data structure generated from ordered minimizer sketches. The tool can be used in a variety of different applications, including improving a draft assembly with a reference-grade genome, a short read assembly with a draft long read assembly, and a draft assembly with an assembly from a closely-related species. When scaffolding a human short read assembly using the reference human genome or a long read assembly, ntJoin improves the NGA50 length 23- and 13-fold, respectively, in under 13 m, using less than 11 GB of RAM. Compared to existing reference-guided assemblers, ntJoin generates highly contiguous assemblies faster and using less memory.

**Availability and implementation:** ntJoin is written in C++ and Python, and is freely available at https://github.com/bcgsc/ntjoin.

## Introduction

Producing highly contiguous assemblies enables important downstream research such as genetic association studies and cis-regulatory element analysis (Rice and Green, 2019). However, while the advancement of single molecule sequencing data such as linked reads and long reads has shown great promise in improving *de novo* genome assembly quality (Shafin, et al., 2019; Weisenfeld, et al., 2017), most draft assemblies are still not achieving chromosome-scale completeness.

For some draft genomes, more contiguous assemblies may be available for a different individual of the same species or even a closely related species. In this case, sequence synteny between the assemblies can be leveraged for assembly-guided scaffolding. For example, while long read assemblies can generate contiguous draft genomes, the high error rates of the reads negatively impacts the base quality, hindering gene annotation (Watson and Warr, 2019). Polishing using short reads is often used to improve the base-pair accuracy of the assemblies (Rice and Green, 2019; Watson and Warr, 2019). An alternative approach to this polishing step is to assemble short reads separately and scaffold the short-read assembly using a long read assembly, producing an assembly on par with the contiguity and structure of the long read assembly, and the base-pair accuracy of the short read assembly.

Existing reference-guided scaffolders such as Ragout (Kolmogorov, et al., 2018) and Ragoo (Alonge, et al., 2019) rely on alignments of the draft assembly to a reference assembly; Ragout utilizes Progressive Cactus (Armstrong, et al., 2019) for large genomes, while Ragoo depends on minimap2 (Li, 2018) for the task. The use of minimizer sketches in tools such as minimap2 is very effective in compactly representing genome sequences. Instead of storing every word of size k (k-mer) from the input sequences, only a chosen set of k-mers or hash values (“minimizers”) are retained, greatly reducing the computational cost of sequence data storage and manipulation (Roberts, et al., 2004).

Here we introduce ntJoin, an assembly-guided scaffolder, which uses a lightweight, alignment-free mapping strategy in lieu of alignments to quickly contiguate a target assembly using one or more references.

## Methods

Given the input target and reference sequence(s), ntJoin first creates an ordered minimizer sketch for each of the supplied sets of sequences, as described previously (Roberts, et al., 2004) (Fig. S1, S2). ntJoin then uses the ordered minimizer sketches from each input to build a single undirected graph that facilitates a lightweight mapping between them. In this graph, each node is a minimizer, and edges are created between minimizers that are adjacent in at least one of the ordered sketches. The graph is then subjected to a series of filtering steps. First, a global edge weight threshold is applied. Next, branching nodes (nodes with degree > 2) are identified, and incident edges are filtered with an increasing edge weight threshold until the degree of that node drops to less than 3. This results in the graph being a set of connected components, each of which is a linear path of minimizer nodes. The sequences of minimizers in the linear paths are then translated to ordered and oriented contig paths, which describe the final output scaffolds. This graph-based method allows the algorithm to perform misassembly correction in addition to scaffolding the input contigs based on the input reference assembly.

Detailed methods are available online.

## Results and Discussion

We first tested ntJoin using various draft and reference-grade *C. elegans* and *H. sapiens* assemblies. Compared to Ragout and Ragoo, ntJoin generally produces assemblies with a higher NGA50 length (length that captures 50% of the genome corrected for misassemblies and genome size), and comparable or fewer misassemblies (Fig. 1; Fig. S4, S6, S10; Tables S3-11). Notably, ntJoin improves assemblies with contiguity in the kilobase range to megabase scale (NGA50 increases from 26.9 kb to 2.3 Mbp and 19.8 kb to 50.3 Mbp, for *C. elegans* and *H. sapiens* short read assemblies, respectively, Fig. S4, S6), while reducing the misassemblies by over a third (33.5% and 61.5%, respectively). This highlights the potential of ntJoin in improving fragmented draft assemblies. Compared to Ragout, ntJoin achieved NGA50 values 1.1-to 2-fold higher for the short read ABySS assemblies tested, although Ragout did scaffold a long read Shasta assembly to a 1.2-fold higher NGA50 (Fig. 1; Fig. S4, S6). However, the Progressive Cactus alignment required for Ragout was very computationally expensive, running for over four days for all human runs and using over 115 GB of RAM, compared to the human ntJoin runs, which finished in under 13 min and used less than 11 GB of RAM. ntJoin was also faster than Ragoo in all tests, from 2.8 times faster for the Shasta assembly up to 71 times faster for the more fragmented *H. sapiens* ABySS assembly (Fig. 1; Fig. S4, S6).

**Fig. 1.**
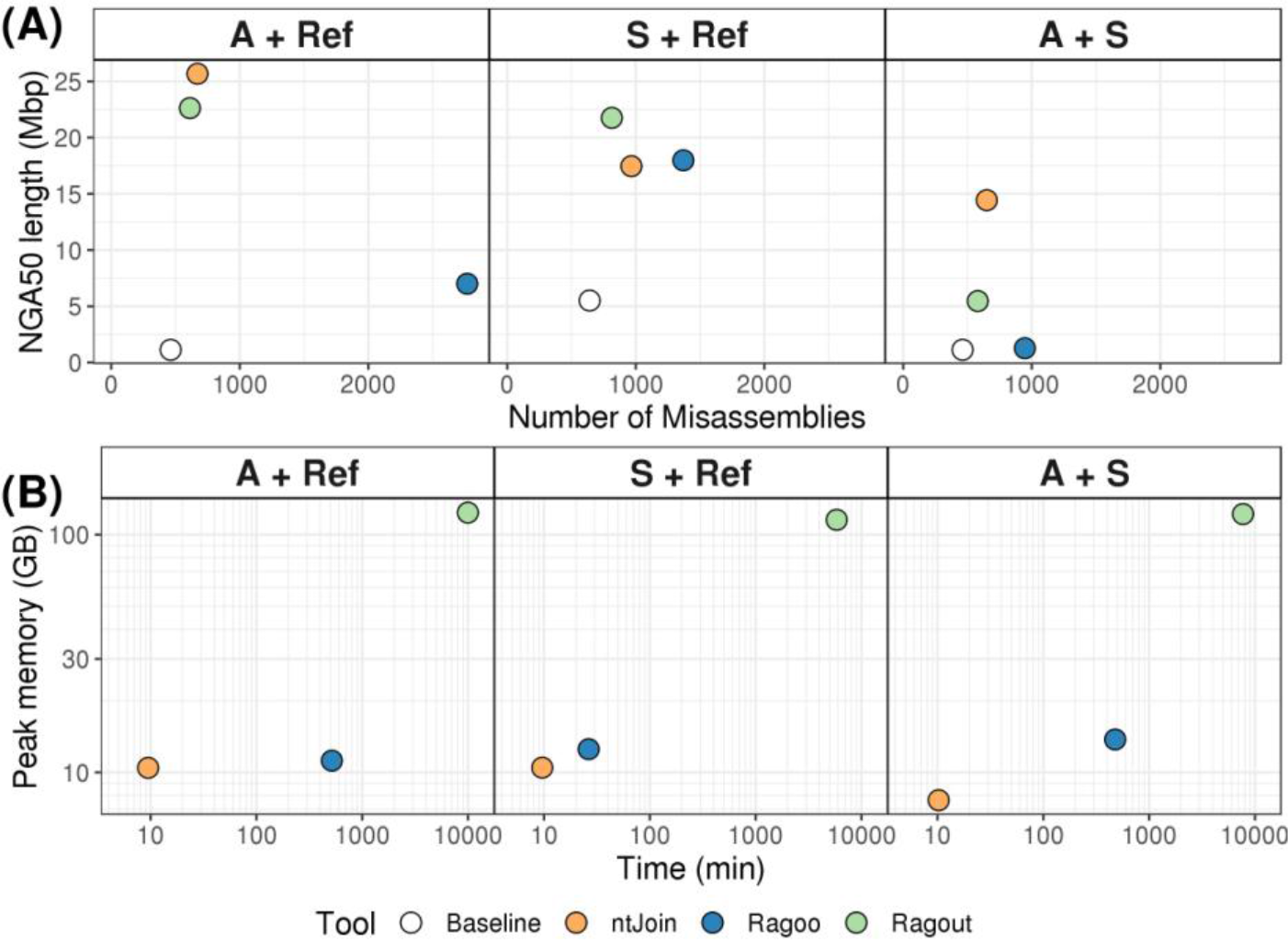
Comparing (A) the contiguity, correctness, and (B) benchmarking results of ntJoin (orange), Ragoo (blue) and Ragout (green) runs on various *H. sapiens* (NA12878) assemblies. The reference genomes are the human reference genome ("Ref") and a polished Shasta assembly ("S"). The draft assemblies improved are a NA12878 ABySS assembly scaffolded with MPET data ("A"), and a polished Shasta assembly ("S").

ntJoin can also improve draft assemblies when contiguous assemblies are available for the same species. In Figure 1, a short read ABySS assembly was scaffolded using a long read Shasta (Shafin, et al., 2019) assembly. By retaining joins unique to the long and short read sequences, ntJoin achieves an NGA50 higher than the baseline Shasta assembly. ntJoin provides an alternative assembly pipeline, where the structure of a long read assembly informs the placement of short read assembly sequences, precluding the need for polishing a long read assembly with short reads. This approach can produce assemblies with high contiguity and base accuracy – particularly important for downstream genome annotation (Tables S12-S15). On this data, neither Ragout nor Ragoo yield assemblies with a similarly high NGA50, and both require more time and memory compared with ntJoin.

The ntJoin approach also extends to scaffolding assemblies of different species, as demonstrated by scaffolding the saltwater and gharial crocodile assemblies using the American alligator genome as reference (Tables S16-S17). In our tests, the NG50 of the crocodile assemblies increased to 14.44 Mbp and 12.92 Mbp for saltwater and gharial crocodiles (baseline NG50 0.14 Mbp and 0.07 Mbp), with a corresponding increase in BUSCO (Simão, et al., 2015) gene completeness of 2.6% and 9.5%, respectively. This demonstrates that ntJoin can still leverage synteny between these target and reference assemblies despite the species having diverged around 80 million years ago (Delsuc, et al., 2018).

In conclusion, ntJoin performs minimizer graph-based scaffolding quickly and with a small memory footprint, while still producing chromosome-level contiguity. As demonstrated, it is a flexible, alignment-free scaffolding tool that can be used in a number of different applications, including hybrid assembly and population genomics research.

