## Supplementary File for "ntJoin: Fast and lightweight assembly-guided scaffolding using minimizer graphs"

### Supplementary Data

#### Supplementary Methods

##### *Generating ordered minimizer sketches*

Minimizer sketches are generated for the input target assembly and reference sequence(s) based on the approach described in Roberts, et al. (2004). Briefly, given a k-mer size (*k*) and a window size (*w*), we compute the hash values of *w* adjacent k-mers using ntHash (Mohamadi, et al., 2016) and select the smallest hash value as the minimizer. This process is repeated for each window of *w* adjacent k-mers in the sequence, producing an ordered sketch of hash values ("minimizers") for each input sequence. As well as storing the ordered minimizers for each input assembly, we also track the contig and position of origin for each minimizer.

##### *Minimizer and graph filtering*

First, the sketches are filtered to only retain minimizers that are unique within a given assembly, and found in each of the input assemblies. These filtered, ordered minimizer sketches are then used to build an undirected graph, where each node is a minimizer, and edges are created between minimizers that are adjacent in at least one of the ordered sketches. The user can provide a weight for each input assembly, and the graph edge weights are the sum of weights of each input assembly that support the edge.

Graph edge filtering occurs in two stages. First, a global edge weight threshold (parameter -*n*, default 1) is applied to the graph. Next, any branch nodes (nodes with degree greater than 2) are identified, and the incident edges filtered with an increasing threshold (from *n* up to max edge weight) until the degree is less than or equal to 2. At this point, the graph is a set of linear minimizer paths which we translate into contig paths for scaffolding.

See Supplementary Figure S1 for a schematic describing the building and filtering of this graph.

##### *Translating minimizer paths to base-pair sequences*

Because we track the source contig and position of the minimizers for each input assembly, the sequence of minimizers in the linear paths can be translated back to contig paths, resulting in ordered and oriented scaffolded contigs (Supplementary Figure S1). Briefly, the linear paths of minimizers are translated to runs of minimizers with the same contig ID in the draft assembly ("contig minimizer runs"). The ordering of the contig minimizer runs defines the order of the contigs in the path. To assign the orientation of a given contig, the positions of all adjacent minimizers in the contig minimizer run are compared to determine if they are largely increasing or decreasing. If the positions are increasing or decreasing at least *m*% of the time (parameter -*m*, default 90), the orientation is assigned as forward or reverse, respectively. If the parameter *mkt=True* is specified, a Mann-Kendall test (Hussain and Mahmud, 2019) is used instead to test if the trend of the positions is significantly increasing or decreasing. This setting is more computationally intensive, especially for short contigs, and we observe that the "majority increasing/decreasing" approach works similarly well. Contigs with an associated contig minimizer run of length one or those that do not satisfy the specified orientation threshold are not assigned an orientation, and will not be incorporated into a path.

We also use the minimizer positions of the reference assembly in the minimizer graph to determine gap sizes (number of 'N's) between scaffolded target contigs. As mentioned previously, the order of contig minimizer runs determines the order of the draft assembly contigs in the paths. These contig minimizer runs are joined together by edges in the graph created based on the reference assembly. For a given pair of contigs merged by ntJoin (*cA, cB*), there is an associated edge between terminal minimizers of these contigs in the minimizer graph (*mA, mB*) (Supplementary Figure S3). Since this edge is connecting two different contigs in the target assembly, these minimizers must be adjacent in a sequence in the reference. Therefore, the estimated gap size is calculated as:

$$gap\ size = |pos(mA, ref) - pos(mB, ref)| - a - b$$

where $a = distance\ from\ pos(mA, draft)\ to\ end\ of\ cA$ and
$b = distance\ from\ pos(mB, draft)\ to\ end\ of\ cB$

Calculating *a, b* in the formula above compensates for the distance from the terminal minimizer in the target contig to the end of the contig, as the terminal minimizers are not always at the very end of the contigs. If multiple reference assemblies are being used, the gap distance is the mean of the estimations from each supporting reference assembly.

##### *Misassembly detection and correction*

As shown in Supplementary Figure S1, if there is a misassembly in the input target contigs (shown in red), the minimizers of a contig may not fully map to the same region of the reference, causing a branching point in the minimizer graph (node with degree > 2). In that case, as long as the reference assembly has a higher user-specified weight than the target assembly, the graph filtering step will remove the edges supported by only the target assembly. This causes different regions of the misassembled contigs to be incorporated into different paths. Thus, ntJoin will cut the input contig at that putative misassembly point, and the regions of the broken contig will then be incorporated into the correct position, as suggested by the minimizer graph. See Supplementary Figure S2 for more details.

##### *Benchmarking scaffolding runs*

To scaffold various draft assemblies using a reference genome, we first assembled paired-end *C. elegans* data using ABySS (v2.1.4; k=64, kc=3, j=8, B=10G, N=9, S=1000-10000) (Jackman, et al., 2017) (Table S2). The ABySS assembly was scaffolded using ntJoin with the *C. elegans* (Bristol N2 strain) reference genome using a sweep of *k* and *w* values (v1.0.1; t=4, target_weight=1, reference_weights='2', n=2) (Table S1, S3).

We then scaffolded three *H. sapiens* (NA12878) assemblies with ntJoin; two short read assemblies and a long read assembly (Table S1- S3). The first short read assembly was generated by assembling 2x250 bp Illumina data using ABySS (v2.1.5; k=128, kc=3, j=25, B=115G, l=40, H=4, S=1000-10000, N=9) (Jackman, et al., 2017). The second short read assembly used the same 2x250 bp Illumina data up to the contig stage of ABySS, but then used 2x101 bp nxtrim-processed MPET reads for the scaffolding stage (O'Connell, et al., 2015). Finally, we ran Shasta (shasta-Linux-0.1.0) on NA12878 release 6 nanopore reads (--input rel_6.fasta) and the resulting assembly polished iteratively using ntEdit (v1.2.2; -i 5 -d 5 -m 1 -t 48 -k 50, 45, 40, 35, 30 ) (Shafin, et al., 2019; Warren, et al., 2019). All human assemblies were scaffolded with ntJoin using the GRCh38 build of the human reference genome (v1.0.1; t=4, target_weight=1, reference_weights='2', n=2) with a sweep of k and w.

We subsequently used ntJoin to scaffold a short read *H. sapiens* assembly using a draft long read *H. sapiens* assembly as the reference. For this test, we used the *H. sapiens* (NA12878) ABySS assembly scaffolded with MPET data as the target assembly and the Shasta assembly with and without polishing as the reference assembly (v1.0.1; t=4, target_weight=1, reference_weights='2', n=1) (Table S1).

To compare the results of ntJoin with existing reference-guided scaffolding tools, we ran each ntJoin scaffolding experiment described above using Ragout (Kolmogorov, et al., 2018) and Ragoo (Alonge, et al., 2019). Ragout was run using default parameters, and the input 'hal' alignment generated with Progressive Cactus (Armstrong, et al., 2019). To generate the alignments in a reasonable time frame, 64 threads were used for the human Progressive Cactus alignments, while four threads were used for the *C. elegans* alignments. Ragoo uses minimap2 (v2.17-r941) (Li, 2018) for the alignment stage, and was run using the chimeric contig detection option (v1.1; -t 4 -b -C). We found that when using a draft assembly as the reference, Ragoo omitted some sequences in the output files. To account for this, we tallied the sequences missing from the output scaffolds file, and rescued those sequences prior to subsequent analysis.

The contiguity and correctness of each assembly was assessed using Quast (v5.0.2; -t 24 --fast --scaffold-gap-max-size 100000 --split-scaffolds --large) (Mikheenko, et al., 2018) using the corresponding reference genome. In comparing the assemblies generated by the various assemblers, we used QUAST-identified misassemblies as well as NGA50, an NG50 contiguity metric adjusted for misassemblies.

In addition to the ntJoin tests using assemblies of the same species, we also used ntJoin to scaffold saltwater (*Crocodylus porosus*) and gharial *(Gavialis gangeticus)* crocodile assemblies using the American alligator (*Alligator mississippiensis*) reference genome (v1.0.1; t=4, target_weight=1, reference_weights='2', k=32, w=500, n=1) (Table S1).

All benchmarking was run using a DELL server with 128 Intel(R) Xeon(R) CPU E7–8867 v3, 2.50GHz with 2.6 TB RAM.

#### Supplementary Figures

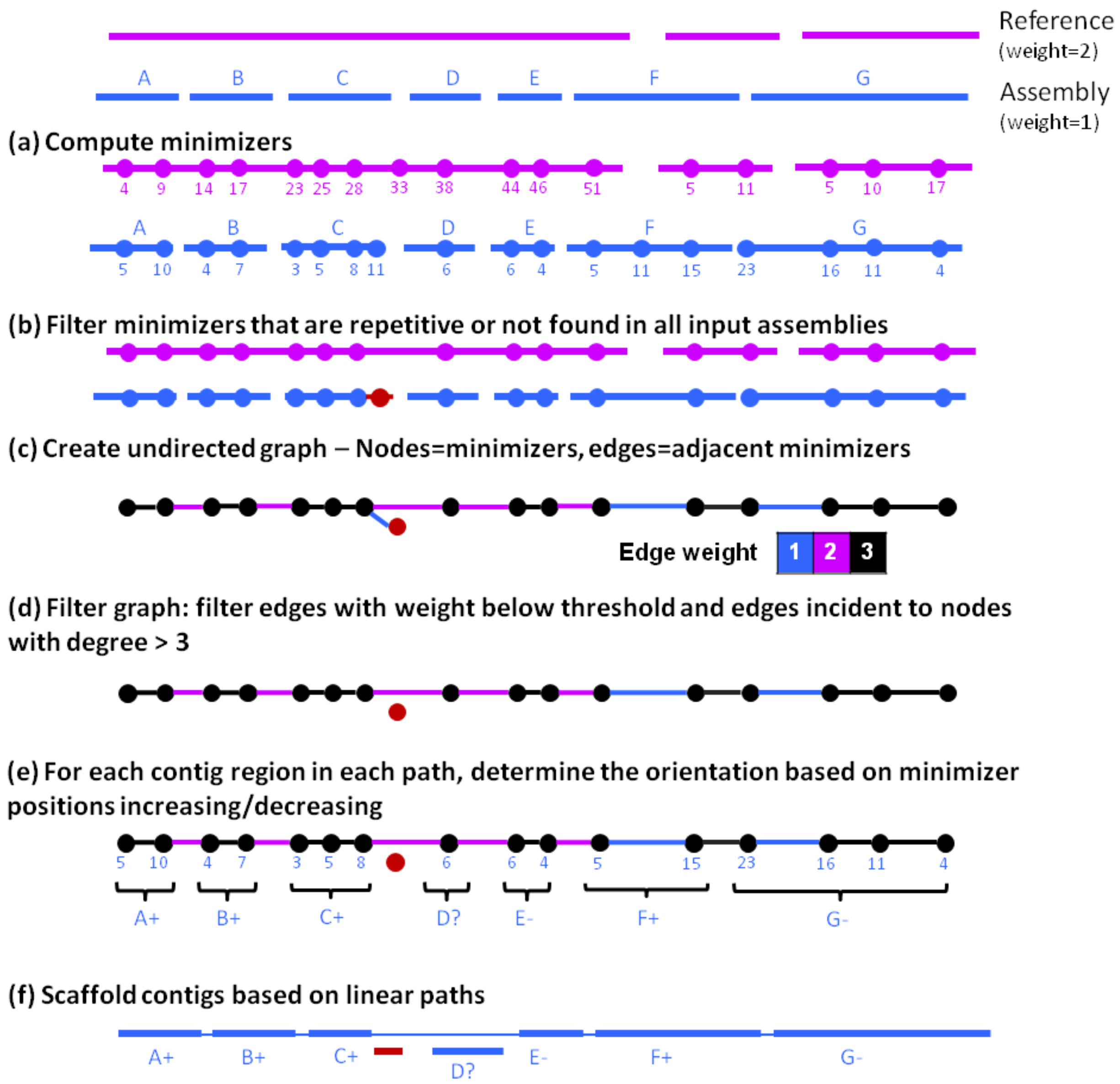


**Supplementary Figure S1. The ntJoin algorithm**. In this example, the target assembly indicated in blue is being scaffolded using the reference assembly, indicated in purple. **(a)** First, ordered minimizer sketches along with their base pair coordinates (indicated as numbers below vertices) are generated for each input assembly. **(b)** Next, minimizers that are repetitive (seen multiple times within the assembly) or unique to an assembly (not found in all input assemblies) are filtered from the sketches. **(c)** The filtered, ordered minimizer sketches are then used to create an undirected minimizer graph, where the nodes are minimizers and edges are created between minimizers that are adjacent in at least one of the assembly sketches. The edge weights are the sum of the weights of the assemblies supporting the edge. For example, in the schematic, if the incident minimizers are only adjacent in the reference, the edge has weight=2, but if the minimizers are adjacent in both the reference and the assembly, the edge has weight=3. **(d)** The graph is first filtered globally, removing edges with a weight less than the specified threshold (*n*, default 1). Next, the edges incident to nodes with a degree > 2 are filtered with an increasing edge weight threshold until the degree is <= 2. **(e)** This graph filtering results in linear paths of minimizers. Because we track the origin contig and position of each minimizer, these minimizer paths can also be seen as a series of runs of minimizers from the same contig (“contig minimizer runs”). The order of the contig minimizer runs defines the contig ordering in the output paths. Whether the run of minimizer positions are largely increasing (positive) or decreasing (negative) is used to infer the relative orientation of each contig. If there is only 1 minimizer for a contig in the path, it cannot be oriented. **(f)** The contigs are scaffolded based on the ordered and oriented contig paths.

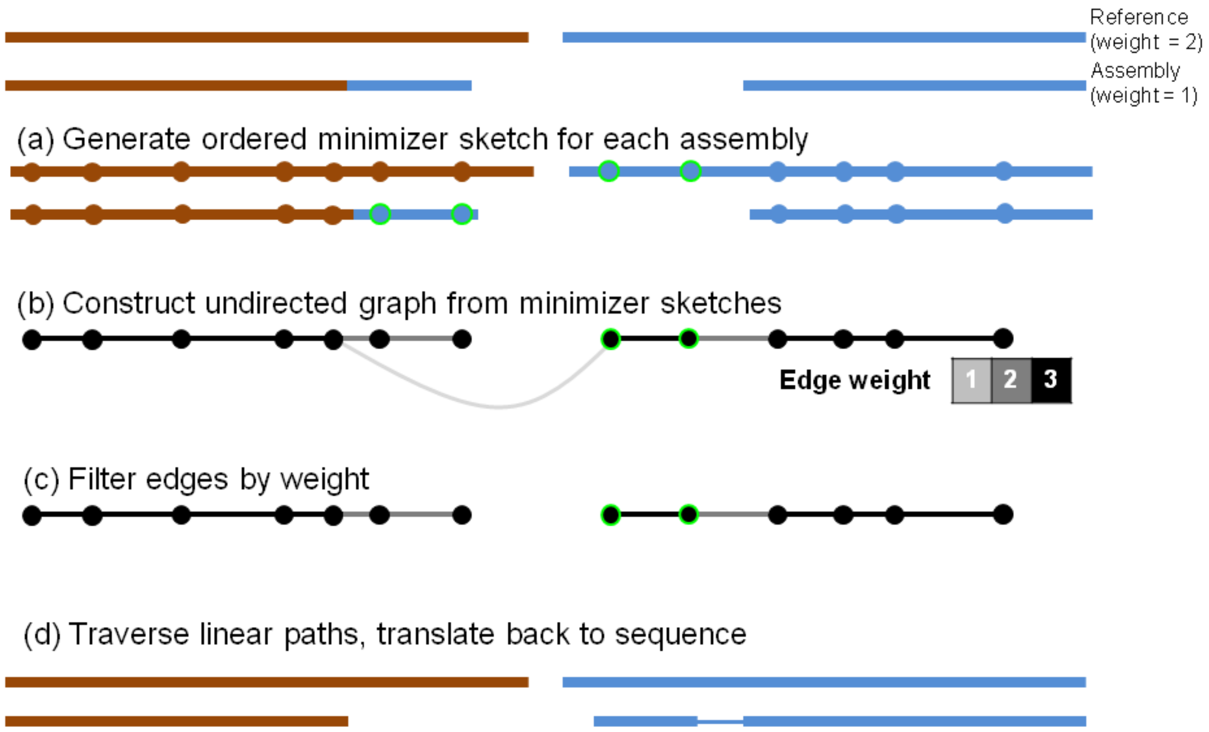


**Supplementary Figure S2**. **Example of misassembly detection and correction in ntJoin.** In this example, the colours (brown, blue) indicate different chromosomes. There is a misassembly in the target assembly, as evident from the chimeric colouring on the first assembly scaffold. **(a, b)** First, ntJoin generates ordered minimizer sketches for both sequences, and constructs an undirected graph where the nodes are minimizers, and edges are created between adjacent minimizers. The minimizers from the misassembled segment of the target assembly are indicated by a green outline. **(c)** The misassembly creates a branch in the undirected graph. The incident edges of that branch node are filtered with an increasing edge weight threshold. For the branch node, the incident edge found only in the target assembly is filtered out (weight = 1), while the incident edge found only in the reference is retained (weight = 2). **(d)** After the edge filtering, two linear paths remain, which are then translated back to sequence, resulting in the correction of the misassembly.

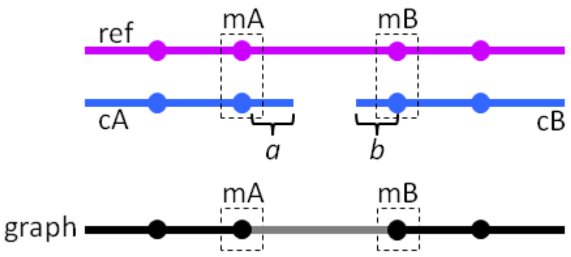


**Supplementary Figure S3. Inferring gap distances in ntJoin**. In this example, contigs *cA* and *cB* in the target assembly with terminal minimizers *mA, mB* are joined based on these minimizers being adjacent in the reference sequence. *a* and *b* are the distances (in bp) between the terminal minimizers of *cA* and *cB* and the ends of the contigs. The gap distance is calculated as: $gap\ size = |pos(mA, ref) - pos(mB, ref)| - a - b$ .

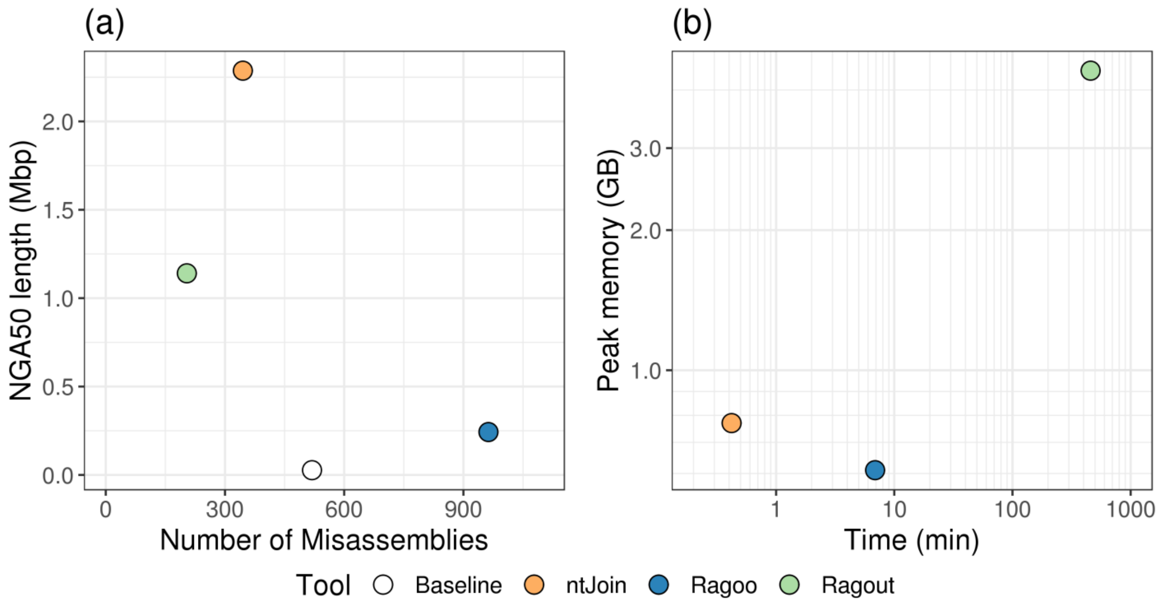


**Supplementary Figure S4.** Correctness, contiguity and benchmarking results of scaffolding an ABySS *C. elegans* short read assembly using the *C. elegans* Bristol N2 reference genome with ntJoin, Ragout and Ragoo.

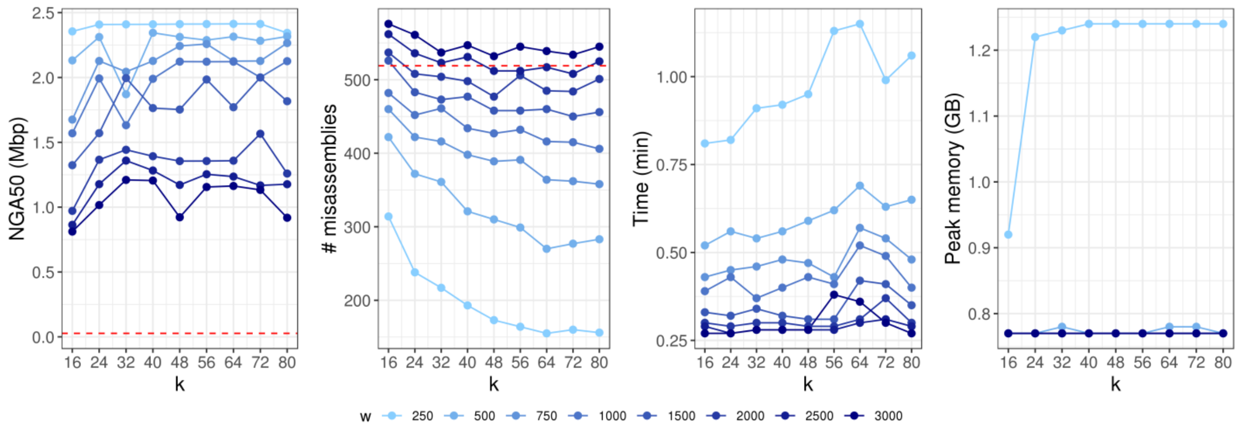


**Supplementary Figure S5.** Scaffolding an ABySS *C. elegans* (Bristol N2) short read assembly using the *C. elegans* Bristol N2 reference genome with ntJoin, sweeping on the k and w parameters. The assembly statistics of the baseline ABySS assembly are indicated by the horizontal dashed red line.

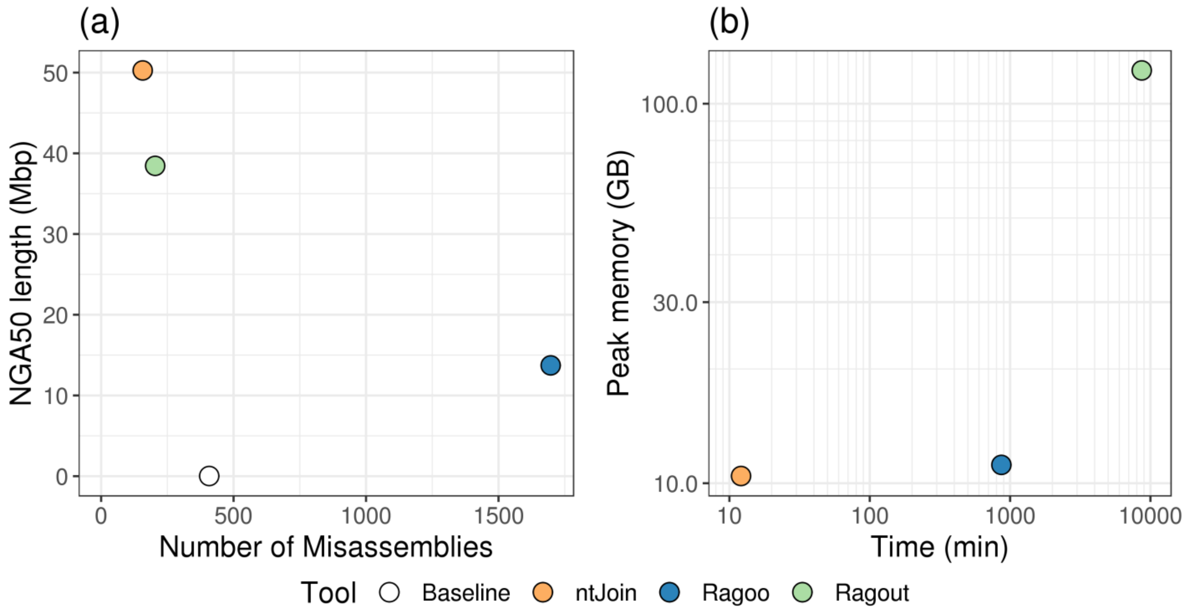


**Supplementary Figure S6.** Correctness, contiguity and benchmarking results of scaffolding an ABySS *H. sapiens* (NA12878) short read assembly using the human reference genome (GRCh38) with ntJoin, Ragout and Ragoo.

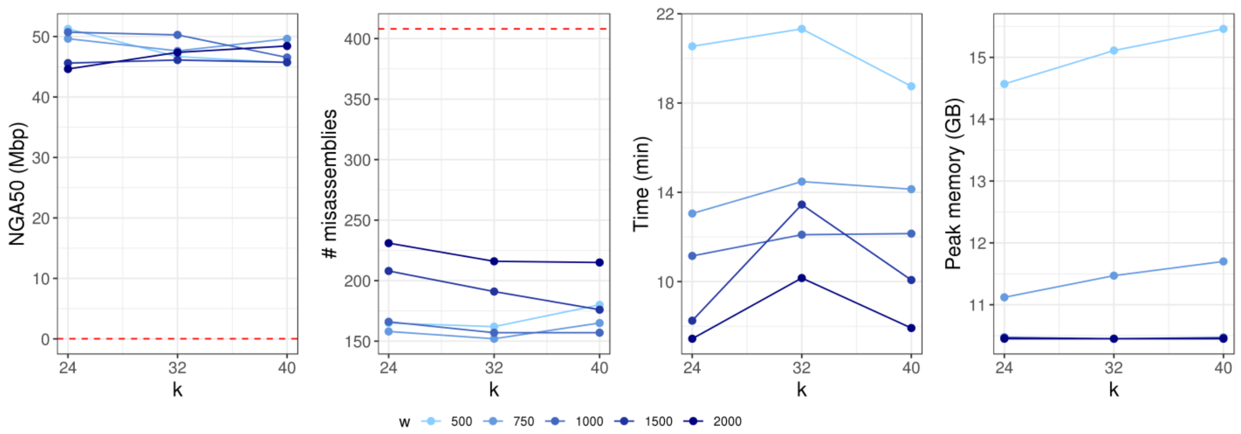


**Supplementary Figure S7.** Scaffolding an ABySS *H. sapiens* (NA12878) short read assembly using the human reference genome (GRCh38) with ntJoin, sweeping on the k and w parameters. The assembly statistics of the baseline ABySS assembly are indicated by the horizontal dashed red line.

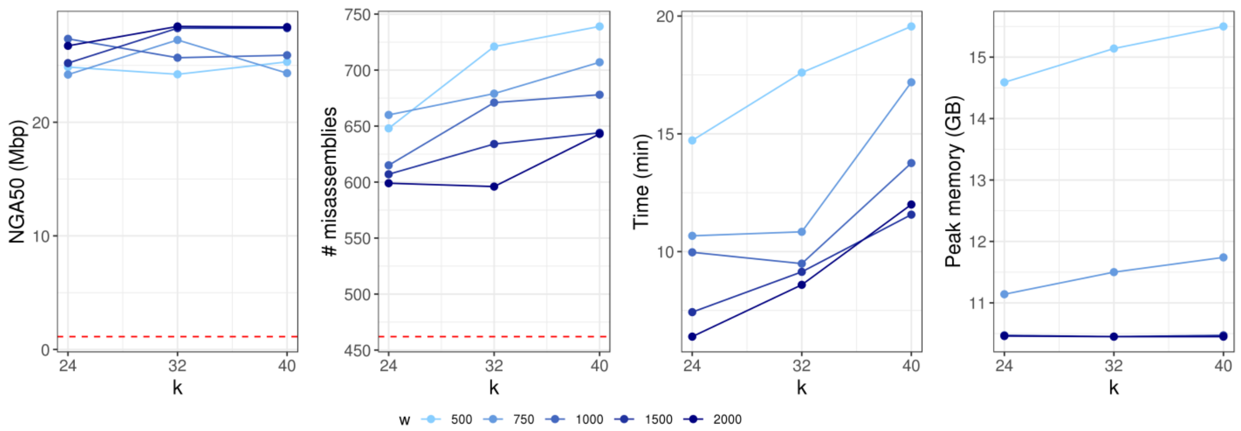


**Supplementary Figure S8.** Scaffolding an ABySS *H. sapiens* (NA12878) short read assembly scaffolded with MPET data using the human reference genome (GRCh38) with ntJoin, sweeping on the k and w parameters. The assembly statistics of the baseline ABySS assembly are indicated by the horizontal dashed red line.

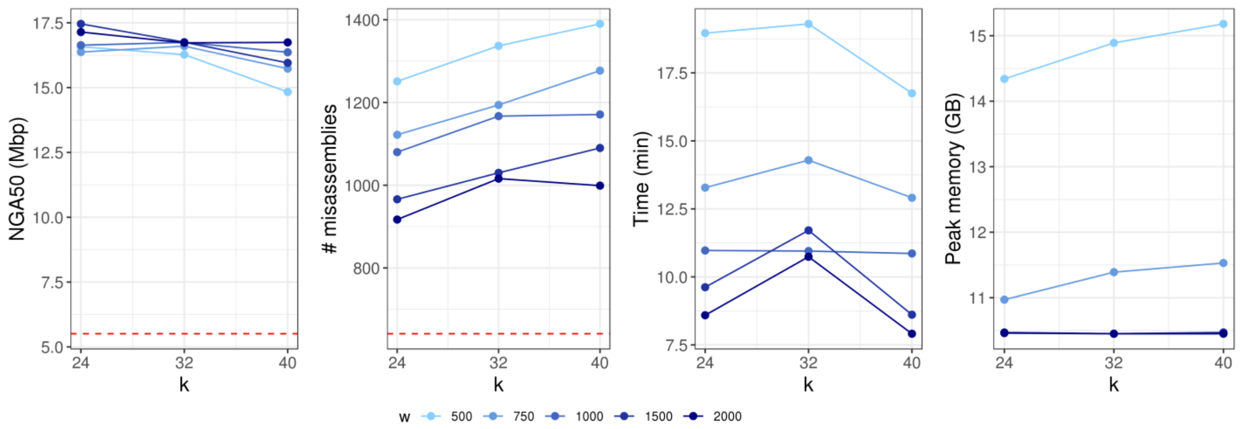


**Supplementary Figure S9.** Scaffolding a Shasta *H. sapiens* (NA12878) long read assembly polished with ntEdit using the human reference genome (GRCh38) with ntJoin, sweeping on the k and w parameters. The assembly statistics of the baseline Shasta assembly are indicated by the horizontal dashed red line.

**Supplementary Figure S10.** Jupiter plot (Chu, 2018) showing consistency between various human genome assemblies and the reference human genome (GRCh38).

**Target: *H. sapiens* (NA12878) ABySS assembly + MPET scaffolding; Reference: *H. sapiens* reference genome GRCh38**

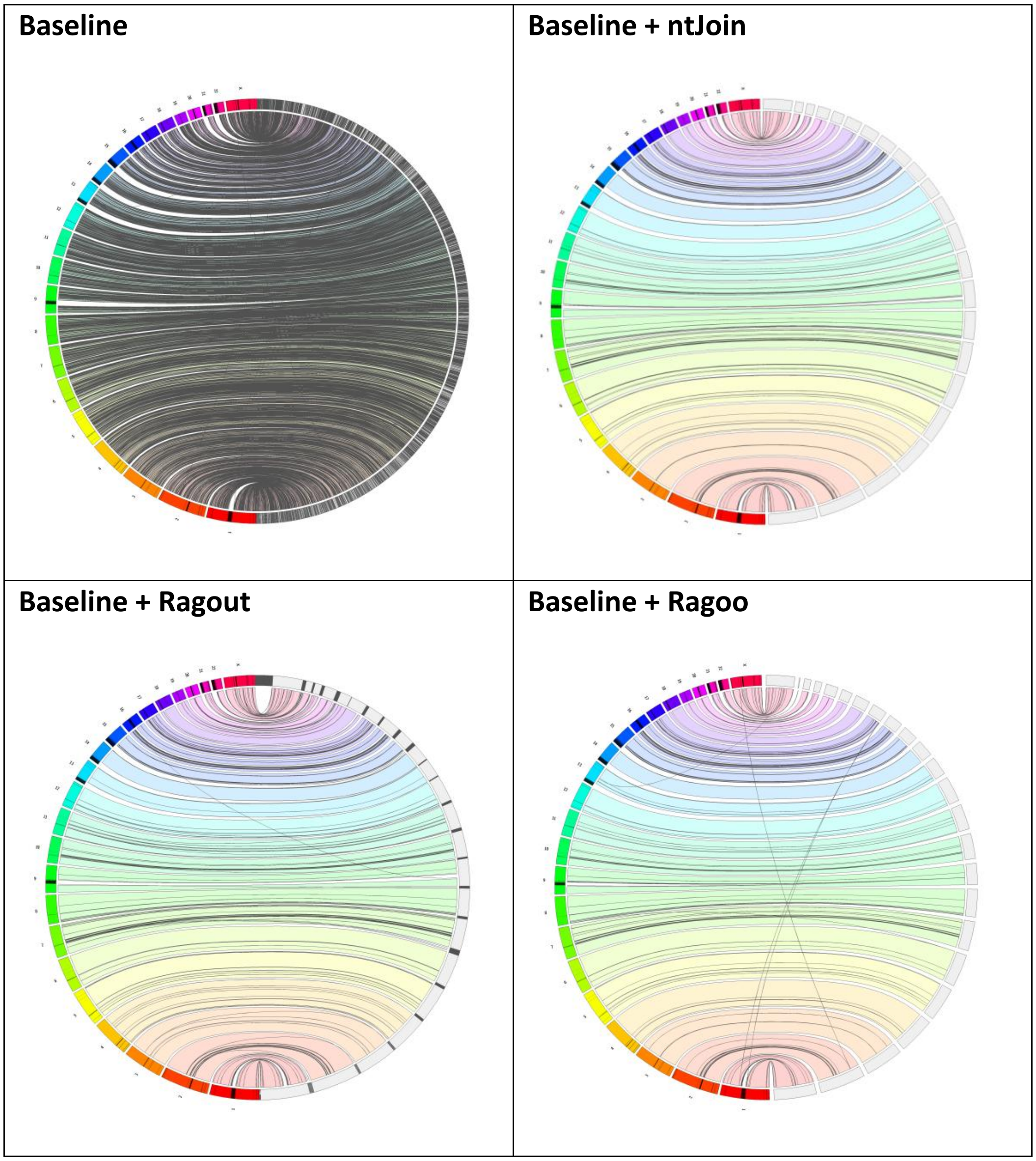

#### Supplementary Tables

**Supplementary Table S1.** Reference and draft assemblies used in our experiments.

| Species | Assembly | URL |
|---|---|---|
| ***C. elegans*** | Bristol N2 reference | https://ftp.ncbi.nlm.nih.gov/genomes/refseq/invertebrate/Caenorhabditis_elegans/reference/GCF_000002985.6_WBcel235/GCF_000002985.6_WBcel235_genomic.fna.gz |
| ***C. elegans*** | ABySS | https://www.bcgsc.ca/downloads/btl/ntJoin/assemblies/celegans/abyss/celegans-abyss.fa |
| ***H. sapiens*** | GRCh8 reference build | https://ftp.ncbi.nlm.nih.gov/genomes/refseq/vertebrate_mammalian/Homo_sapiens/latest_assembly_versions/GCF_000001405.39_GRCh38.p13/GCF_000001405.39_GRCh38.p13_genomic.fna.gz |
| ***H. sapiens*** | ABySS | https://www.bcgsc.ca/downloads/btl/ntJoin/assemblies/hsapiens/NA12878_abyss/hsapiens-abyss.fa |
| ***H. sapiens*** | ABySS (+ MPET) | https://www.bcgsc.ca/downloads/btl/ntJoin/assemblies/hsapiens/NA12878_abyss-mpet/hsapiens-abyss-mpet.fa |
| ***H. sapiens*** | Shasta | https://www.bcgsc.ca/downloads/btl/ntJoin/assemblies/hsapiens/NA12878_shasta/hsapiens-shasta.fa |
| ***H. sapiens*** | Shasta + ntEdit | https://www.bcgsc.ca/downloads/btl/ntJoin/assemblies/hsapiens/NA12878_shasta-ntedit/hsapiens-shasta-ntedit.fa |
| ***Crocodylus porosus*** | NCBI accession GCA_000768395.1 | ftp://ftp.ncbi.nlm.nih.gov/genomes/all/GCA/000/768/395/GCA_000768395.1_Cpor_2.0/GCA_000768395.1_Cpor_2.0_genomic.fna.gz |
| ***Gavialis gangeticus*** | NCBI accession GCA_000775435.1 | ftp://ftp.ncbi.nlm.nih.gov/genomes/all/GCA/000/775/435/GCA_000775435.1_ggan_v0.2/GCA_000775435.1_ggan_v0.2_genomic.fna.gz |
| ***Alligator mississippiensis*** | NCBI accession GCA_000281125.4 | ftp://ftp.ncbi.nlm.nih.gov/genomes/all/GCA/000/281/125/GCA_000281125.4_ASM28112v4/GCA_000281125.4_ASM28112v4_genomic.fna.gz |

**Supplementary Table S2.** Sequencing reads used for draft assemblies.

| Species | Reads | URL |
|---|---|---|
| ***C. elegans* N2** | Illumina (2x110 bp) | ftp://ftp.sra.ebi.ac.uk/vol1/fastq/DRR008/DRR008444 |
| ***H. sapiens* NA12878** | Illumina (2x250 bp) | https://basespace.illumina.com/ ("HiSeq 2500: TruSeq PCR-Free DNA 2x251 (NA12878)") |
| ***H. sapiens* NA12878** | Illumina mate-pair (2x101 bp) | ftp://ftp.ddbj.nig.ac.jp/ddbj_database/dra/fastq/ERA207/ERA207860/ERX237515/ERR262997.fastq.bz2 |

**Supplementary Table S3.** Contiguity and correctness statistics of baseline assemblies used for ntJoin, Ragout and Ragoo runs.

| Species | Assembly | Scaffolds >= 3kb | NG50 (bp) | NGA50 (bp) | Misassemblies | Mismatches per 100kb | Indels per 100kb |
|---|---|---|---|---|---|---|---|
| *C. elegans* | ABySS | 7,891 | 30,293 | 26,876 | 519 | 13.42 | 4.96 |
| *H. sapiens* | ABySS | 160,260 | 19,900 | 19,794 | 408 | 93.81 | 24.06 |
| *H. sapiens* | ABySS + MPET | 6,656 | 1,175,297 | 1,131,267 | 462 | 120.81 | 28.07 |
| *H. sapiens* | Shasta | 2,410 | 6,917,709 | 5,414,750 | 603 | 224.58 | 525.30 |
| *H. sapiens* | Shasta + ntEdit | 2,411 | 7,119,313 | 5,507,381 | 641 | 230.87 | 103.30 |

**Supplementary Table S4.** Benchmarking results of scaffolding a *C. elegans* ABySS short read assembly with ntJoin using the *C. elegans* Bristol N2 reference genome. The assembly indicated in bold is discussed in the main text.

| k | w | Scaffolds >= 3kb | NG50 (Mbp) | NGA50 (Mbp) | Misassemblies | Time (min) | Peak Memory (GB) |
|---|---|---|---|---|---|---|---|
| 16 | 250 | 1,005 | 17.58 | 2.36 | 314 | 0.64 | 0.91 |
| 16 | 500 | 2,166 | 17.56 | 2.13 | 422 | 0.37 | 0.77 |
| 16 | 750 | 2,846 | 17.52 | 1.68 | 460 | 0.33 | 0.77 |
| 16 | 1,000 | 3,185 | 17.51 | 1.57 | 482 | 0.35 | 0.77 |
| 16 | 1,500 | 3,695 | 17.50 | 1.32 | 526 | 0.25 | 0.77 |
| 16 | 2,000 | 4,109 | 17.48 | 0.97 | 537 | 0.22 | 0.77 |
| 24 | 250 | 534 | 17.56 | 2.41 | 238 | 0.69 | 1.22 |
| 24 | 500 | 1,747 | 17.55 | 2.31 | 372 | 0.41 | 0.77 |
| 24 | 750 | 2,516 | 17.52 | 2.13 | 422 | 0.31 | 0.77 |
| 24 | 1,000 | 2,845 | 17.51 | 1.99 | 452 | 0.30 | 0.77 |
| 24 | 1,500 | 3,344 | 17.51 | 1.57 | 483 | 0.23 | 0.77 |
| 24 | 2,000 | 3,793 | 17.50 | 1.37 | 508 | 0.23 | 0.77 |
| 32 | 250 | 464 | 17.57 | 2.41 | 217 | 0.68 | 1.23 |
| **32** | **500** | **1,669** | **17.53** | **2.29** | **345** | **0.42** | **0.77** |
| 32 | 750 | 2,502 | 17.53 | 2.04 | 416 | 0.35 | 0.77 |
| 32 | 1,000 | 2,811 | 17.52 | 2.01 | 444 | 0.26 | 0.77 |
| 32 | 1,500 | 3,356 | 17.50 | 2.00 | 473 | 0.26 | 0.77 |
| 32 | 2,000 | 3,756 | 17.49 | 1.44 | 504 | 0.21 | 0.77 |
| 40 | 250 | 384 | 17.59 | 2.41 | 193 | 0.73 | 1.24 |
| 40 | 500 | 1,628 | 17.57 | 2.34 | 321 | 0.41 | 0.77 |
| 40 | 750 | 2,463 | 17.52 | 2.13 | 398 | 0.34 | 0.77 |
| 40 | 1,000 | 2,814 | 17.52 | 1.99 | 434 | 0.27 | 0.77 |
| 40 | 1,500 | 3,327 | 17.51 | 1.76 | 477 | 0.26 | 0.77 |
| 40 | 2,000 | 3,747 | 17.50 | 1.39 | 498 | 0.22 | 0.77 |
| 48 | 250 | 349 | 17.58 | 2.41 | 173 | 0.71 | 1.24 |
| 48 | 500 | 1,609 | 17.57 | 2.31 | 310 | 0.42 | 0.77 |
| 48 | 750 | 2,463 | 17.56 | 2.24 | 389 | 0.33 | 0.77 |
| 48 | 1,000 | 2,788 | 17.51 | 2.12 | 427 | 0.31 | 0.77 |
| 48 | 1,500 | 3,323 | 17.50 | 1.75 | 458 | 0.23 | 0.77 |
| 48 | 2,000 | 3,726 | 17.49 | 1.36 | 477 | 0.22 | 0.77 |
| 56 | 250 | 334 | 17.58 | 2.41 | 164 | 0.70 | 1.24 |
| 56 | 500 | 1,601 | 17.56 | 2.29 | 299 | 0.40 | 0.77 |
| 56 | 750 | 2,485 | 17.56 | 2.26 | 391 | 0.32 | 0.77 |
| 56 | 1,000 | 2,815 | 17.55 | 2.12 | 432 | 0.32 | 0.77 |
| 56 | 1,500 | 3,308 | 17.51 | 1.98 | 458 | 0.24 | 0.77 |
| 56 | 2,000 | 3,729 | 17.50 | 1.36 | 506 | 0.23 | 0.77 |
| 64 | 250 | 325 | 17.59 | 2.41 | 155 | 0.76 | 1.24 |
| 64 | 500 | 1,586 | 17.57 | 2.32 | 270 | 0.41 | 0.78 |
| 64 | 750 | 2,435 | 17.53 | 2.13 | 364 | 0.33 | 0.77 |
| 64 | 1,000 | 2,775 | 17.56 | 2.12 | 416 | 0.30 | 0.77 |
| 64 | 1,500 | 3,297 | 17.54 | 1.77 | 460 | 0.25 | 0.77 |
| 64 | 2,000 | 3,744 | 17.49 | 1.36 | 485 | 0.24 | 0.77 |

**Supplementary Table S5.** Benchmarking results of scaffolding a *C. elegans* ABySS short read assembly with Ragout and Ragoo using the *C. elegans* Bristol N2 reference genome.

| Tool | Scaffolds >= 3kb | NG50 (Mbp) | NGA50 (Mbp) | Misassemblies | Time (min) | Peak memory (GB) |
|---|---|---|---|---|---|---|
| Ragout | 1,365 | 17.49 | 1.14 | 204 | 458.41 | 4.40 |
| Ragoo | 2,054 | 16.26 | 0.24 | 963 | 6.88 | 0.61 |

**Supplementary Table S6.** Benchmarking results of scaffolding a human (NA12878) ABySS short read assembly with ntJoin using the human reference genome GRCh38. The assembly indicated in bold is discussed in the main text.

| k | w | Scaffolds >= 3kb | NG50 (Mbp) | NGA50 (Mbp) | Misassemblies | Time (min) | Peak Memory (GB) |
|---|---|---|---|---|---|---|---|
| 24 | 500 | 1,804 | 153.37 | 51.26 | 165 | 20.54 | 14.57 |
| 24 | 750 | 2,239 | 153.24 | 49.64 | 158 | 13.05 | 11.12 |
| 24 | 1,000 | 3,063 | 153.24 | 50.72 | 166 | 11.15 | 10.47 |
| 24 | 1,500 | 6,212 | 145.34 | 45.60 | 208 | 8.25 | 10.47 |
| 24 | 2,000 | 12,558 | 153.18 | 44.63 | 231 | 7.44 | 10.45 |
| 32 | 500 | 1,569 | 153.41 | 46.68 | 162 | 21.32 | 15.11 |
| 32 | 750 | 1,866 | 151.89 | 47.62 | 152 | 14.48 | 11.47 |
| **32** | **1,000** | **2,370** | **145.41** | **50.27** | **157** | **12.10** | **10.45** |
| 32 | 1,500 | 4,797 | 145.35 | 46.12 | 191 | 13.45 | 10.45 |
| 32 | 2,000 | 10,787 | 145.31 | 47.38 | 216 | 10.16 | 10.45 |
| 40 | 500 | 1,441 | 153.41 | 45.67 | 180 | 18.75 | 15.46 |
| 40 | 750 | 1,649 | 145.45 | 49.62 | 165 | 14.14 | 11.70 |
| 40 | 1,000 | 2,011 | 145.41 | 46.53 | 157 | 12.15 | 10.47 |
| 40 | 1,500 | 3,910 | 151.78 | 45.74 | 176 | 10.07 | 10.46 |
| 40 | 2,000 | 9,443 | 153.18 | 48.43 | 215 | 7.92 | 10.45 |

**Supplementary Table S7.** Benchmarking results of scaffolding a human (NA12878) ABySS short read assembly with Ragout and Ragoo using the human reference genome.

| Tool | Scaffolds >= 3kb | NG50 (Mbp) | NGA50 (Mbp) | Misassemblies | Time (min) | Peak memory (GB) |
|---|---|---|---|---|---|---|
| Ragout | 6,443 | 153.25 | 38.45 | 204 | 8,669.67 | 122.36 |
| Ragoo | 94 | 138.08 | 13.72 | 1,697 | 866.63 | 11.18 |

**Supplementary Table S8.** Benchmarking results of scaffolding a human (NA12878) ABySS short read assembly (scaffolded with MPET data) with ntJoin using the human reference genome GRCh38. The assembly shown in Figure 1 is bolded.

| k | w | Scaffolds >= 3kb | NG50 (Mbp) | NGA50 (Mbp) | Misassemblies | Time (min) | Peak Memory (GB) |
|---|---|---|---|---|---|---|---|
| 24 | 500 | 1,326 | 143.11 | 24.86 | 648 | 14.72 | 14.59 |
| 24 | 750 | 1,340 | 140.74 | 24.19 | 660 | 10.67 | 11.14 |
| 24 | 1,000 | 1,364 | 140.59 | 27.34 | 615 | 9.97 | 10.47 |
| 24 | 1,500 | 1,541 | 142.28 | 25.20 | 607 | 7.43 | 10.47 |
| 24 | 2,000 | 1,691 | 142.45 | 26.72 | 599 | 6.39 | 10.46 |
| 32 | 500 | 1,161 | 143.04 | 24.22 | 721 | 17.60 | 15.14 |
| 32 | 750 | 1,212 | 142.73 | 27.24 | 679 | 10.84 | 11.50 |
| **32** | **1,000** | **1,237** | **141.39** | **25.68** | **671** | **9.49** | **10.45** |
| 32 | 1,500 | 1,410 | 140.67 | 28.28 | 634 | 9.14 | 10.45 |
| 32 | 2,000 | 1,586 | 142.44 | 28.43 | 596 | 8.59 | 10.45 |
| 40 | 500 | 1,075 | 142.88 | 25.31 | 739 | 19.56 | 15.50 |
| 40 | 750 | 1,134 | 142.74 | 24.32 | 707 | 17.19 | 11.74 |
| 40 | 1,000 | 1,135 | 142.67 | 25.90 | 678 | 13.76 | 10.47 |
| 40 | 1,500 | 1,297 | 142.69 | 28.28 | 644 | 11.57 | 10.47 |
| 40 | 2,000 | 1,465 | 140.66 | 28.38 | 643 | 12.00 | 10.45 |

**Supplementary Table S9.** Benchmarking results of scaffolding a human ABySS (NA12878) short read assembly (scaffolded with MPET data) with Ragout and Ragoo using the human reference genome GRCh38.

| Tool | Scaffolds >= 3kb | NG50 (Mbp) | NGA50 (Mbp) | Misassemblies | Time (min) | Peak memory (GB) |
|---|---|---|---|---|---|---|
| Ragout | 1,851 | 141.81 | 22.62 | 612 | 9,971.82 | 123.73 |
| Ragoo | 47 | 138.21 | 7.01 | 2,769 | 517.07 | 11.20 |

**Supplementary Table S10.** Benchmarking results of scaffolding a human (NA12878) Shasta long read assembly (polished with ntEdit) with ntJoin using the human reference genome GRCh38. The assembly shown in Figure 1 is bolded.

| k | w | Scaffolds >= 3kb | NG50 (Mbp) | NGA50 (Mbp) | Misassemblies | Time (min) | Peak Memory (GB) |
|---|---|---|---|---|---|---|---|
| 24 | 500 | 950 | 154.66 | 16.58 | 1,251 | 18.96 | 14.34 |
| 24 | 750 | 1,005 | 155.59 | 16.37 | 1,122 | 13.28 | 10.97 |
| 24 | 1,000 | 1,058 | 150.06 | 16.64 | 1,080 | 10.97 | 10.47 |
| **24** | **1,500** | **1,130** | **145.04** | **17.46** | **966** | **9.62** | **10.47** |
| 24 | 2,000 | 1,180 | 152.53 | 17.14 | 917 | 8.59 | 10.46 |
| 32 | 500 | 902 | 156.82 | 16.27 | 1,337 | 19.30 | 14.89 |
| 32 | 750 | 960 | 150.65 | 16.60 | 1,194 | 14.29 | 11.39 |
| 32 | 1,000 | 994 | 155.42 | 16.74 | 1,167 | 10.95 | 10.45 |
| 32 | 1,500 | 1,053 | 142.12 | 16.75 | 1,030 | 11.71 | 10.45 |
| 32 | 2,000 | 1,121 | 142.26 | 16.73 | 1,016 | 10.74 | 10.45 |
| 40 | 500 | 868 | 146.56 | 14.83 | 1,390 | 16.75 | 15.18 |
| 40 | 750 | 927 | 146.05 | 15.73 | 1,277 | 12.91 | 11.53 |
| 40 | 1,000 | 985 | 151.74 | 16.36 | 1,171 | 10.86 | 10.47 |
| 40 | 1,500 | 1,063 | 142.59 | 15.95 | 1,090 | 8.61 | 10.47 |
| 40 | 2,000 | 1,107 | 142.01 | 16.74 | 999 | 7.91 | 10.45 |

**Supplementary Table S11.** Benchmarking results of scaffolding a human (NA12878) Shasta long read assembly (polished with ntEdit) with Ragout and Ragoo using the human reference genome GRCh38.

| Tool | Scaffolds >= 3kb | NG50 (Mbp) | NGA50 (Mbp) | Misassemblies | Time (min) | Peak memory (GB) |
|---|---|---|---|---|---|---|
| Ragout | 1,350 | 142.28 | 21.75 | 814 | 5,833.22 | 115.36 |
| Ragoo | 105 | 140.88 | 17.98 | 1,370 | 26.42 | 12.52 |

**Supplementary Table S12.** Benchmarking results of scaffolding a human (NA12878) ABySS short read assembly (scaffolded with MPET data) with ntJoin using an unpolished Shasta long read assembly as reference.

| k | w | Scaffolds >= 3kb | NG50 (Mbp) | NGA50 (Mbp) | Misassemblies | Mismatches per 100 kbp | Indels per 100 kbp | Time (min) | Peak Memory (GB) |
|---|---|---|---|---|---|---|---|---|---|
| 24 | 500 | 2,779 | 16.61 | 10.99 | 929 | 117.36 | 28.29 | 15.95 | 14.93 |
| 24 | 750 | 2,567 | 17.77 | 11.79 | 786 | 119.56 | 28.21 | 13.19 | 10.99 |
| 24 | 1,000 | 2,375 | 20.75 | 13.27 | 715 | 120.46 | 28.17 | 10.22 | 8.85 |
| 24 | 1,500 | 2,437 | 20.54 | 13.57 | 639 | 120.88 | 28.12 | 8.75 | 6.90 |
| 24 | 2,000 | 2,460 | 22.59 | 13.93 | 612 | 120.91 | 28.09 | 7.36 | 5.81 |
| 32 | 500 | 2,762 | 19.51 | 11.63 | 893 | 117.12 | 28.36 | 16.55 | 15.09 |
| 32 | 750 | 2,543 | 18.94 | 12.89 | 762 | 119.63 | 28.21 | 14.47 | 11.09 |
| 32 | 1,000 | 2,379 | 21.35 | 13.51 | 722 | 120.3 | 28.15 | 11.77 | 8.89 |
| 32 | 1,500 | 2,408 | 22.58 | 14.00 | 665 | 120.81 | 28.11 | 9.03 | 6.92 |
| 32 | 2,000 | 2,460 | 23.23 | 13.63 | 606 | 120.76 | 28.08 | 7.77 | 5.80 |
| 40 | 500 | 2,710 | 16.11 | 10.72 | 913 | 117.16 | 28.41 | 12.48 | 15.08 |
| 40 | 750 | 2,480 | 19.17 | 12.75 | 791 | 119.57 | 28.18 | 10.27 | 11.07 |
| 40 | 1,000 | 2,319 | 19.04 | 12.98 | 739 | 120.81 | 28.17 | 7.94 | 8.90 |
| 40 | 1,500 | 2,322 | 19.77 | 13.94 | 667 | 120.6 | 28.09 | 6.12 | 6.92 |
| 40 | 2,000 | 2,463 | 18.10 | 13.57 | 633 | 121.02 | 28.1 | 5.71 | 5.82 |

**Supplementary Table S13.** Benchmarking results of scaffolding a human (NA12878) ABySS short read assembly (scaffolded with MPET data) with Ragout and Ragoo using an unpolished Shasta long read assembly as reference.

| Tool | Scaffolds >= 3kb | NG50 (Mbp) | NGA50 (Mbp) | Misassemblies | Mismatches per 100 kbp | Indels per 100 kbp | Time (min) | Peak memory (GB) |
|---|---|---|---|---|---|---|---|---|
| Ragout | 2,975 | 7.12 | 5.42 | 598 | 111.61 | 29.35 | 7,546.63 | 146.86 |
| Ragoo | 5,260 | 0.97 | 0.90 | 906 | 122.42 | 27.93 | 480.87 | 13.77 |

**Supplementary Table S14.** Benchmarking results of scaffolding a human (NA12878) ABySS short read assembly (scaffolded with MPET data) with ntJoin using an ntEdit-polished Shasta long read assembly as reference. The assembly shown in Figure 1 is bolded.

| k | w | Scaffolds >= 3kb | NG50 (Mbp) | NGA50 (Mbp) | Misassemblies | Mismatches per 100 kbp | Indels per 100 kbp | Time (min) | Peak Memory (GB) |
|---|---|---|---|---|---|---|---|---|---|
| 24 | 500 | 2,646 | 17.68 | 12.93 | 880 | 116.49 | 28.24 | 17.31 | 16.34 |
| 24 | 750 | 2,438 | 18.68 | 12.89 | 736 | 119.32 | 28.21 | 14.34 | 12.08 |
| 24 | 1,000 | 2,276 | 22.58 | 13.65 | 695 | 120.38 | 28.19 | 10.80 | 9.49 |
| 24 | 1,500 | 2,334 | 23.43 | 13.93 | 643 | 120.68 | 28.1 | 8.88 | 7.42 |
| 24 | 2,000 | 2,342 | 23.31 | 14.12 | 650 | 120.82 | 28.1 | 7.04 | 6.15 |
| 32 | 500 | 2,567 | 19.66 | 12.64 | 894 | 115.22 | 28.33 | 18.10 | 17.01 |
| 32 | 750 | 2,357 | 20.56 | 13.16 | 775 | 118.77 | 28.2 | 14.92 | 12.51 |
| 32 | 1,000 | 2,207 | 21.12 | 13.51 | 749 | 120.04 | 28.14 | 11.31 | 9.81 |
| **32** | **1,500** | **2,258** | **23.36** | **14.43** | **650** | **120.58** | **28.13** | **10.27** | **7.64** |
| 32 | 2,000 | 2,328 | 23.23 | 14.01 | 620 | 120.6 | 28.1 | 9.40 | 6.28 |
| 40 | 500 | 2,531 | 19.97 | 13.05 | 968 | 114.79 | 28.36 | 14.38 | 17.45 |
| 40 | 750 | 2,335 | 22.60 | 12.91 | 813 | 119.47 | 28.19 | 11.55 | 12.80 |
| 40 | 1,000 | 2,173 | 22.58 | 13.63 | 753 | 120.19 | 28.11 | 8.96 | 10.03 |
| 40 | 1,500 | 2,165 | 21.13 | 13.84 | 651 | 120.58 | 28.11 | 6.99 | 7.80 |
| 40 | 2,000 | 2,254 | 23.24 | 14.28 | 626 | 120.97 | 28.1 | 6.33 | 6.42 |

**Supplementary Table S15.** Benchmarking results of scaffolding a human (NA12878) ABySS short read assembly (scaffolded with MPET data) with Ragout and Ragoo using an ntEdit-polished Shasta long read assembly as reference.

| Tool | Scaffolds >= 3kb | NG50 (Mbp) | NGA50 (Mbp) | Misassemblies | Mismatches per 100 kbp | Indels per 100 kbp | Time (min) | Peak memory (GB) |
|---|---|---|---|---|---|---|---|---|
| Ragout | 2,969 | 6.98 | 5.46 | 580 | 111.66 | 29.35 | 7,718.75 | 122.05 |
| Ragoo | 5,578 | 1.38 | 1.27 | 946 | 120.71 | 28.08 | 477.63 | 13.75 |

**Supplementary Table S16.** Assembly statistics of baseline genome assemblies of *Crocodylus porosus* (saltwater crocodile), *Gavialis gangeticus* (gharial crocodile) and *Alligator mississippiensis* (American alligator). The NG50 length metric is based on an estimated genome size of 2.7 Gbp (St John, et al., 2012), and the “tetrapoda” lineage was used for the BUSCO analysis (Simão, et al., 2015).

| Species | Scaffolds >= 3kb | NG50 (Mbp) | Complete BUSCOs |
|---|---|---|---|
| ***Alligator mississippiensis*** | 3,392 | 12.53 | 3,729 (94.41%) |
| ***Crocodylus porosus*** | 20,596 | 0.14 | 3,520 (89.11%) |
| ***Gavialis gangeticus*** | 38,644 | 0.07 | 3,251 (82.30%) |

**Supplementary Table S17.** Scaffolding the genome assemblies of *Crocodylus porosus* (saltwater crocodile) and *Gavialis gangeticus* (gharial crocodile) with ntJoin (k=32 w=500 n=1) using an *Alligator mississippiensis* (American alligator) genome assembly. The NG50 length metric is based on an estimated genome size of 2.7 Gbp (St John, et al., 2012), and the "tetrapoda" lineage was used for the BUSCO analysis (Simão, et al., 2015).

| Crocodile species | Scaffolds >= 3kb | NG50 (Mbp) | Complete BUSCOs | Time (min) | Peak Memory (GB) |
|---|---|---|---|---|---|
| ***Crocodylus porosus*** | 10, 279 | 14.44 | 3,624 (91.75%) | 3.9 | 7.04 |
| ***Gavialis gangeticus*** | 16,499 | 12.92 | 3,627 (91.82%) | 4.18 | 7.22 |
